# Differential responses of primary neuron-secreted MCP-1 and IL-9 to type 2 diabetes and Alzheimer’s disease-associated metabolites

**DOI:** 10.1101/2023.11.17.567595

**Authors:** Brendan K. Ball, Madison K. Kuhn, Rebecca M. Fleeman, Elizabeth A. Proctor, Douglas K. Brubaker

## Abstract

Type 2 diabetes (T2D) is implicated as a risk factor for Alzheimer’s disease (AD), the most common form of dementia. In this work, we investigated neuroinflammatory responses of primary neurons to potentially circulating, blood-brain barrier (BBB) permeable metabolites associated with AD, T2D, or both. We identified nine metabolites associated with protective or detrimental properties of AD and T2D in literature (lauric acid, asparagine, fructose, arachidonic acid, aminoadipic acid, sorbitol, retinol, tryptophan, niacinamide) and stimulated primary mouse neuron cultures with each metabolite before quantifying cytokine secretion via Luminex. We employed unsupervised clustering, inferential statistics, and partial least squares discriminant analysis to identify relationships between cytokine concentration and disease-associations of metabolites. We identified MCP-1, a cytokine associated with monocyte recruitment, as differentially abundant between neurons stimulated by metabolites associated with protective and detrimental properties of AD and T2D. We also identified IL-9, a cytokine that promotes mast cell growth, to be differentially associated with T2D. Indeed, cytokines, such as MCP-1 and IL-9, released from neurons in response to BBB-permeable metabolites associated with T2D may contribute to AD development by downstream effects of neuroinflammation.

## INTRODUCTION

Alzheimer’s disease (AD) is a neurodegenerative disease characterized by the progressive loss of memory and cognitive impairment, which affects more than 6.7 million people in the United States^1^. There is not a single cause for AD, but rather a multitude of risk factors that are likely responsible, including age, genetics, family history, and pre-existing health conditions^2–4^. Substantial evidence suggests that people with type 2 diabetes (T2D), a chronic metabolic disease that affects the body’s ability to regulate and process glucose, increase the risk for AD^5,6^. While dependent on environmental location and other lifestyle factors, up to 81% of people who have AD have T2D or impaired glucose levels^5,7^. Accounting for age, a shared risk factor in both diseases, T2D is a significant risk factor for the development of AD^8^. The incurred risk of AD from altered glucose metabolism has been demonstrated in both rodent and human studies^9–11^. Some studies pointed to the link between disease co-morbidity to impairment of insulin receptors, which is associated with decreased brain glucose metabolism^12,13^. Other reports demonstrated that chronic inflammation from diabetes results in altered levels of proinflammatory cytokines in the brain, which may serve as a risk factor for AD pathology^14–16^. Despite the clear association between the risk of AD progression with a history of T2D, the biological mechanisms in which T2D pathobiology promotes AD development are not well understood.

The blood-brain barrier (BBB) is a structure of tightly connected brain endothelial cells, pericytes, and astrocytes that protects the brain from harmful substances and ensures the passage of nutrients from circulation into the brain^17,18^. The BBB also regulates the entry of immune cells into the central nervous system and the export of toxic metabolic waste from the brain^19–21^. In some disease states, BBB integrity may become impaired, with the breakdown of tight junctions observed in both human AD and T2D^18,22,23^. When the BBB is impaired, substances that may not typically be transported to the brain, such as circulating metabolites, are more likely to cross over and stimulate local neuronal cells^24^. When stimulated by external factors such as metabolites, neurons and glia cells of the central nervous system may release cytokines to signal proinflammatory or anti-inflammatory responses^25,26^. Additionally, the secretion of proinflammatory cytokines often results in the activation of immune cells, which may lead to the release of additional cytokines and initiate an inflammatory signaling cascade^27^. Subsequently, to return to a homeostatic state, anti-inflammatory cytokines relay messages to reduce or stop the pro-inflammatory response.

Under normal physiological conditions, leukocytes do not readily traffic across the BBB^28^. With cytokines being released by neurons and glia such as astrocytes and microglia, the impairment of the BBB may lead to increased trafficking of unwanted substances and peripheral immune cells such as macrophages and neutrophils into the brain^19,29^. Inflammatory cytokines may impair the BBB and increase the selective membrane’s permeability through the destruction of endothelial tight junctions^30–32^. In this case, the barrier integrity may become further compromised such that peripheral immune cells can enter areas with neuroinflammation^33,34^. Thus, in a condition such as T2D-associated AD development, metabolites or other small molecules originating elsewhere in the body and circulating in the blood may cross the BBB to stimulate the cells in the brain. This chronic, low-grade stimulation of neuronal cells may lead to downstream neuroinflammation, promoting the development of AD^35–39^. However, the potential of systemic circulating species already upregulated in T2D to promote neuroinflammation in the brain has been understudied.

In this work, we identified patterns of cytokine production in primary mouse neurons following stimulation by metabolites differentially produced in T2D and AD. We find that primary neurons differentially secrete cytokines in response to different metabolites based on associations to AD, T2D, or both. Collectively, our findings indicate that disease-associated metabolites and their interactions with neurons may serve an important role in the neuroinflammatory pathway and the potential biological pathway in which T2D increases the risk of AD development.

## RESULTS

### Retinol and Arachidonic Acid Trigger Distinct Cytokine Response Compared to Other Metabolites

We first cultured mouse-derived primary neurons for stimulation. After the neuronal plating, nine different metabolites, which include lauric acid, asparagine, fructose, arachidonic acid, aminoadipic acid, D-sorbitol, retinol, L-tryptophan, and niacinamide was used to induce cytokine secretion by the primary culture of neurons. The cytokines secreted in the neuronal media were then collected and quantified using the Luminex assay (**Fig. 1**).

**Figure 1.**
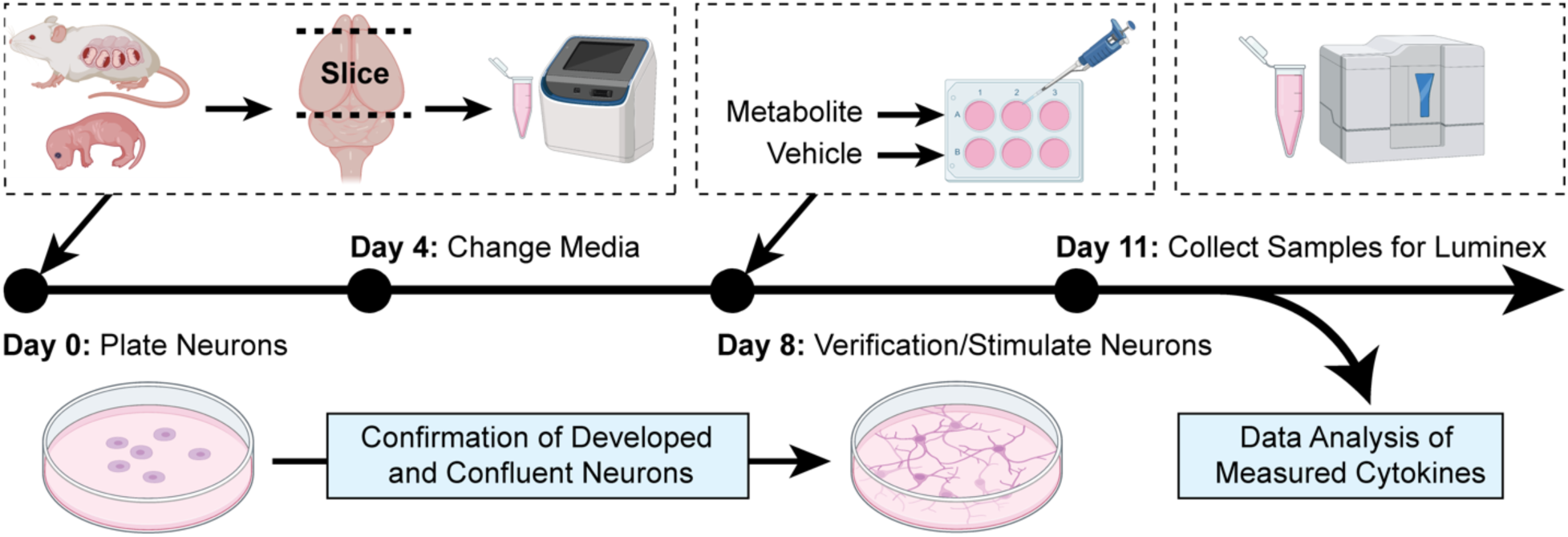
Method in plating and culturing primary neurons derived from embryonic CD1 mice. The embryonic litters from pregnant CD-1 mice were decapitated, and the cortical regions were isolated for primary neuron culturing. On day 8, the metabolites and respective vehicles were used to stimulate the primary neurons. After three days, the neuron media samples were collected for the quantification of released cytokines using the Luminex platform. (Created with BioRender.com).

Of the 23 cytokines quantified, eight had more than 25% of the measurements below the lower limit of quantification (G-CSF, IL-17A, IL-13, IL-2, IL-3, IL-4, IL-5, and TNF-α), and were excluded from the analysis. The data for the fifteen remaining cytokines were pre-processed by replacing quantified values that were below the lowest limit of quantification with the lowest respective standard value (2.1% of data). The cytokine concentrations were normalized by a log_2_ ratio of the cytokine measurement divided by the average of the triplicate vehicle concentrations.

We analyzed the normalized log_2_ fold change of the cytokines with unsupervised clustering methods to discover patterns in the data independent of the experimental conditions. Here, we found that MCP-1, a chemoattractant for monocytes that enhances the recruitment of peripheral immune cells, was upregulated by retinol, L-tryptophan, and niacinamide, all of which are AD-protective metabolites^40^. All metabolites associated with T2D or AD downregulated MCP-1 (**Fig. 2a**). We found that retinol, which is associated with AD/T2D-protective properties, tended to induce larger magnitude fold changes in the cytokines compared to other metabolites. Nine cytokines showed statistically significant differences in abundance between the disease-association groupings of metabolites (Kruskal-Wallis test, FDR *q* value < 0.10), including IL-6, MIP-1β, IL-12p70, IL-9, RANTES, IL-12p40, MCP-1, IL-1β, and IL-10 (**Fig. 2**, **Supplementary Fig. S1**).

**Figure 2.**
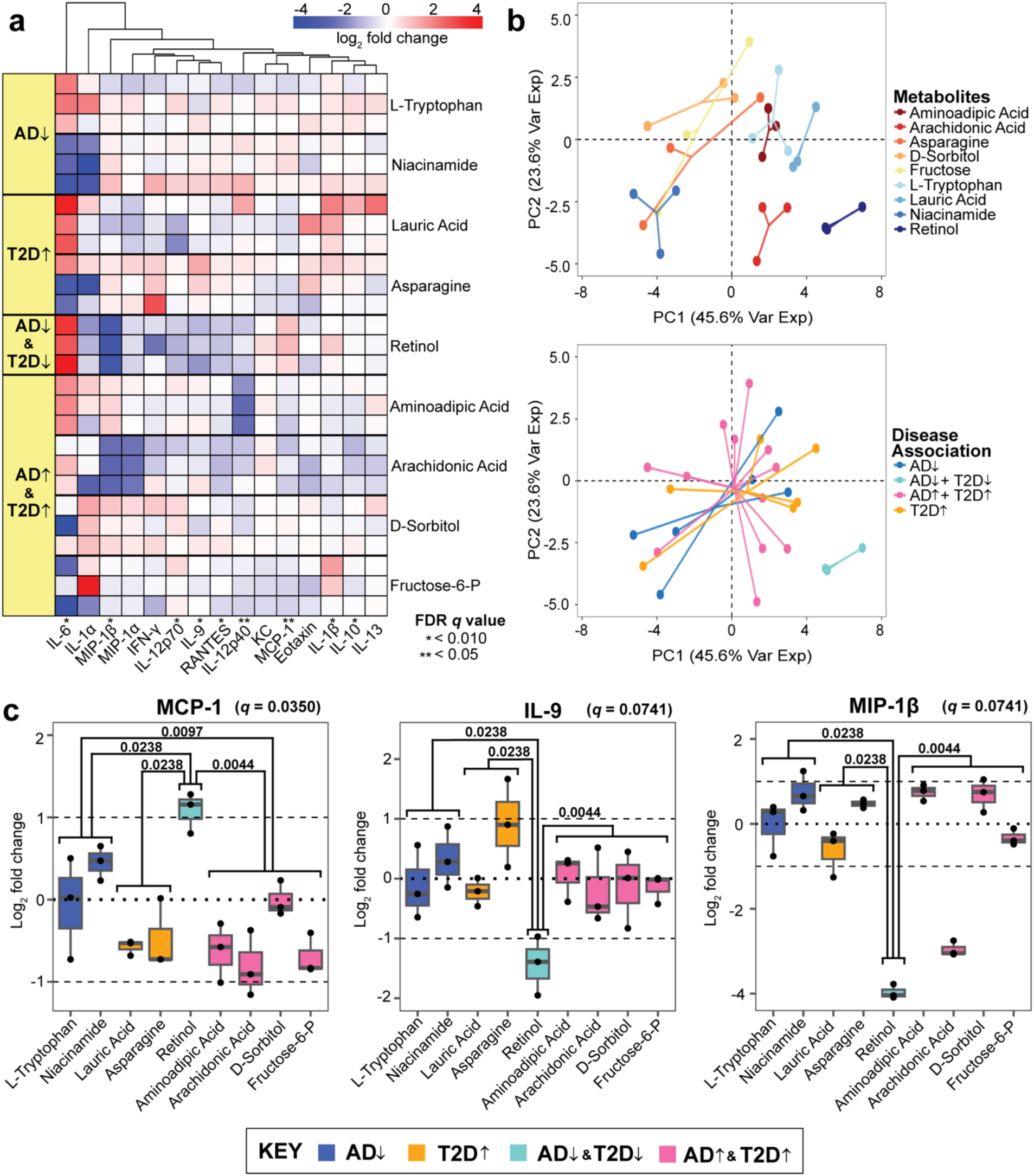
Unsupervised hierarchical clustering of cytokines released by primary neurons. **(a)** Hierarchical clustering of the log_2_ fold change of cytokine concentrations across the nine tested metabolites. **(b)** Principal component analysis of the log_2_ fold changes categorized by individual metabolites and disease-associations. The direction of the arrows indicates disease (↓) and protective (↑) characteristics related to AD and T2D. Each metabolite is associated with a category of disease. Significance was determined from a Kruskal-Wallis test corrected by the Benjamini-Hochberg method (FDR *q* value < 0.10). **(c)** The log_2_ ratio of cytokine concentration to vehicles of MCP-1, IL-9, and MIP-1β. Mann-Whitney pair-wise testing was applied to each metabolite group based on disease association (significance in *p* value denoted within the plot).

We performed a principal component analysis (PCA) to determine which metabolites induced a different neuroinflammatory signaling response on the primary mouse neurons. We found that seven of the nine metabolites, excluding retinol and arachidonic acid, elicited cytokine responses that clustered together regardless of disease association (**Fig. 2b**). This indicates that retinol and arachidonic acid stimulation results in distinct neuronal cytokine responses compared to the other metabolites.

Retinol in particular induced significant downregulation of IL-9 and MIP-1β and up-regulation of MCP-1 in a pattern that deviated significantly from all other cytokine responses to other metabolites (**Fig. 2c**). MCP-1 is a monocyte attractive factor and has functional similarities to other noteworthy cytokine responses in IL-9, and MIP-1β. IL-9 is responsible for a large variety of physiological processes, including the promotion of mast cell growth and function, similar to MCP-1^41^. Likewise, MIP-1β is a chemoattractant that activates human granulocytes, which can lead to acute inflammation^42^. These three cytokines contain at least one significant difference between response to protective and detrimental associations of metabolites but are not consistent with MCP-1. The up-regulated of MCP-1 by retinol in the AD^↓^T2D^↓^ (protective) group, but not the AD-protective group alone, indicates that the retinol-MCP-1 response is specific to the T2D-AD axis and not AD alone.

### Multivariate Statistical Modeling Identifies MCP-1 and IL-9 as T2D Differentiating Cytokines for AD Development

We next assessed whether a supervised modeling approach would identify combinations of cytokines capable of stratifying the metabolites based on different disease association groups. Using a multivariate statistical modeling framework: Partial Least Squares Discriminant Analysis (PLS-DA), we identified cytokines that are most predictive of either protective or disease properties of the disease association of interest. These metabolites were categorized with their respective disease associations: AD and T2D protective (AD^↓^T2D^↓^), AD and T2D (AD^↑^T2D^↑^), AD protective (AD^↓^), and T2D (T2D^↑^). The model first was to identify patterns of cytokine secretion differentiating the groups between AD-protective, AD/T2D-protective, T2D, and AD/T2D, answering the question if certain cytokines are more differentially produced in the presence of specific disease-associated metabolites. With metabolites being complex biological molecules, we next asked if our findings from the 4-way model were consistent if categorized by associations with AD alone or T2D alone. Here, we prepared two other models that separate the metabolites into AD- and T2D-only groups. For the latter two models, any metabolites that did not fit the criteria (e.g., T2D-only metabolite for the AD model) were excluded before applying the cross-validation and constructing the PLS-DA model.

In the 4-way PLS-DA model, we found that the cytokines contributing to the overall PLS-DA model more than average (VIP>1) on both LV1 and LV2 were MCP-1, IL-12p40, and IL-9. (**Fig. 3a-b**). A variable with a VIP>1 in both LV1 and LV2 suggests that the specific cytokine is consistently contributing to the separation across different components of the PLS-DA model. This is important because the identified cytokine is reliable for discriminating between the disease categories. We also found that the AD^↓^T2D^↓^ group separated from the other three other metabolite groups. This indicated that the T2D-AD protective metabolite, retinol, induced a distinct cytokine response while the other three groups, AD-protective, T2D-associated, and AD/T2D-associated, were more similar to each other.

**Figure 3.**
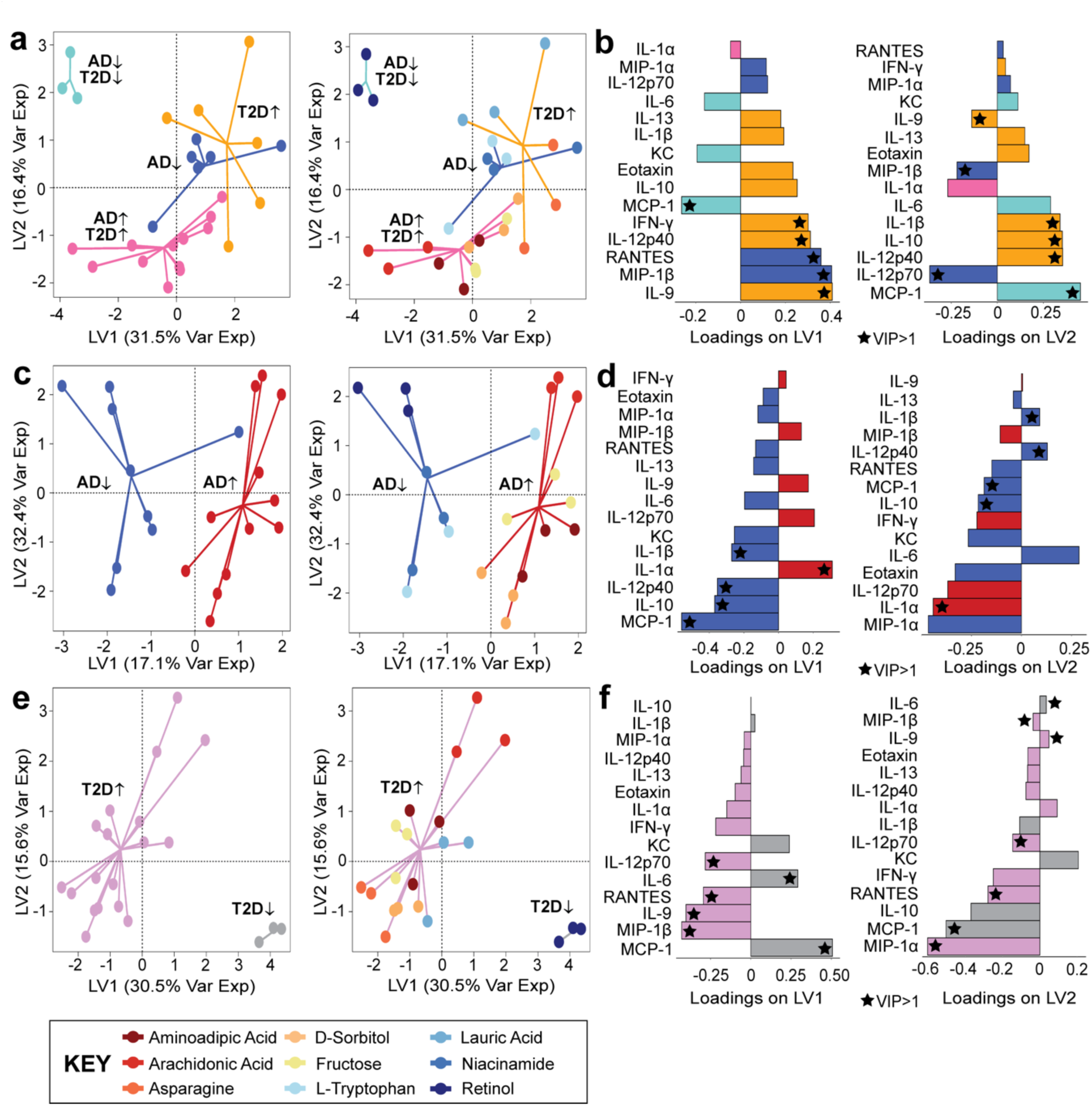
Separation of different disease-associated metabolites is detected from the PLS-DA model. **(a-b)** Four-way disease associations**; (c-d)** AD-protective and AD associated classifications, and **(e-f)** T2D-protective and T2D associated classifications. Loading variables for each model (LV1 and LV2) with a VIP>1 is labeled with a star, and the color of the loading bar represents the cytokine with the highest contribution to the specific class (metabolite grouping).

Having demonstrated a separation among the four-way disease-association classification **(Fig. 3a-b**), we adjusted the model to compare the metabolites based on their association to specific relationships, such as AD/AD-protective or T2D/T2D-protective. We adjusted our model to determine if our findings from the four-way model were consistent across the AD/AD-protective and T2D/T2D-protective models. In our AD-only model (**Fig. 3c-d**), the model separated between the AD disease-associated metabolites and the AD-protective metabolites across LV1. Cytokines with VIP>1 contributing to both LV1 and LV2 were IL-1β, IL-12p40, IL-10, IL-1α, and MCP-1. Similar to the AD PLS-DA model, our T2D PLS-DA model successfully separated the metabolites associated with the protective and detrimental properties of T2D (**Fig. 3e-f**). In our T2D model, six cytokines had a VIP>1 contributing to LV1 and LV2 (IL-12p70, RANTES, IL-6, IL-9, MIP-1β, and MCP-1).

Of the several cytokines contributing a VIP>1 to each of the models, we identified two cytokines that were consistent across each of the three PLS-DA models. The two cytokines were MCP-1 and IL-9. MCP-1 was found to be contributing to the protective properties of T2D and AD, while IL-9 was identified to be contributing to T2D properties. Overall, these three PLS-DA models demonstrated that primary mouse neurons stimulated by metabolites with different disease associations lead to distinct responses by neuroinflammatory cytokines.

## DISCUSSION

Here, we used an *in vitro* primary neuron culture approach to test our hypothesis that BBB-permeable metabolites with AD or T2D associations may stimulate neuroinflammatory or protective processes in neurons associated with disease. Independent research groups have shown that dysregulation of the BBB is a known characteristic shared in both AD and T2D and others have investigated the relationship between cytokine levels and disease status in separate AD and T2D studies^22,23,43,44^. Breakdown of the BBB may serve as an important connection between AD and T2D by allowing circulating factors to cross the BBB and induce disease-associated signaling cascades or provoke local inflammatory responses^35–39^.

Our work shows that MCP-1 is responsive to the AD/T2D protective metabolite retinol. Upregulation of MCP-1 corresponded to protective properties of AD and T2D, whereas downregulation of MCP-1 from primary neurons was a response to disease-associated metabolite stimulation. MCP-1 contributed more than average to each of the three PLS-DA models and MCP-1 was found to be responsive to metabolites associated with protective characteristics. Previous studies that investigated the relationship between serum retinol and AD reported diminished circulating retinol as an increased risk factor for cognitive decline in humans^45^ and in mice^46^. Additionally, retinol was found to be decreased in subjects with T2D, as well as subjects with T2D and diabetic retinopathy compared to healthy groups^47^. This case may be attributed to a derivative of retinol, retinoic acid, which is converted through a two-step oxidation process. Retinoic acid has been demonstrated to restore the insulin function of β-cells in mouse models^48^. Β-cells are localized in the pancreas and are responsible for creating insulin to regulate blood sugar levels. Further understanding the possible protective role of retinol may serve as a viable preventative avenue for T2D-driven AD.

Through literature, we find evidence supporting that a deficiency of MCP-1-activated immune cells, such as microglia, has an increased risk of early onset of AD. We hypothesize that neuron-secreted MCP-1 activates microglia to promote the clearance of amyloid-beta proteins in the brain and, based on our results, retinol stimulation could promote this process. This finding is promising because present in people with AD, amyloid-beta, and tau tangles are the hallmark proteins found aggregated in the brain^49,50^.

Despite our findings, other sources that have studied AD signaling found MCP-1 to be associated with longitudinal cognitive decline in patients with AD^51,52^. However, the MCP-1 cytokine from these studies quantified cytokines from plasma samples, rather than neuron-derived media. Thus, we acknowledge that there may be conflicting cytokine findings from human AD studies^51–54^. This difference in results may be attributed to several factors, such as the high dimensionality of human biology and the location in the body to which the cytokines were quantified.

IL-9 was responsive tod T2D^↑^ associated metabolites (lauric acid and asparagine) and significantly differentially abundant compared to retinol than the AD^↓^T2D^↓^ metabolite. Lauric acid was reported to induce hepatic insulin resistance through the upregulation of SELENOP, which encodes for selenoprotein P^55^. In the case of asparagine, the loss of homeostasis between two interconvertible amino acids (asparagine and aspartate) was found associated with T2D^56^. For both of these metabolites, PLS-DA also revealed IL-9 to be contributing more than average to the classification of T2D. A study by Mohammed *et al*., found that serum IL-9 in T2D patients was higher than that of the healthy control group^57^. A separate study found that lower serum IL-9 was associated with pre-diabetes and T2D^58^. The conflicting results are based on cytokine quantification from serum samples and may not be reflective of the responses we observed in neuronal media. Though IL-9 was not significantly responsive to the specific AD-associated metabolites we tested, others found that higher neuron and astrocyte production of IL-9 *in vitro* is linked to AD progression^59^ and this is also seen in human studies^60^.

Based on our findings, we hypothesize a potentially shared neuroinflammatory response pathway where, during a healthy state (**Fig. 4a**), the BBB regulates the passage of nutrients and substances across the membrane with a tightly locked layer of brain endothelial cells. As a result, upregulated metabolites due to metabolic disease development will have a difficult time crossing the BBB to stimulate the neuronal cells. This ultimately prevents peripheral immune cells from also entering the brain area which will generate a larger neuroinflammatory response. During a diseased state, the BBB breaks down, and harmful substances and unwanted cells can leak across the membrane (**Fig. 4b**). As a result, neurons and other similar cells become stimulated and release cytokines^61^. These cytokines can be pro-inflammatory or anti-inflammatory and trigger other downstream effects that may lead to a positive feedback loop of cytokine secretions. This long-term cycle may ultimately contribute to the disruption of the BBB, as well as chronic neuroinflammation we may observe in people living with AD today.

**Figure 4.**
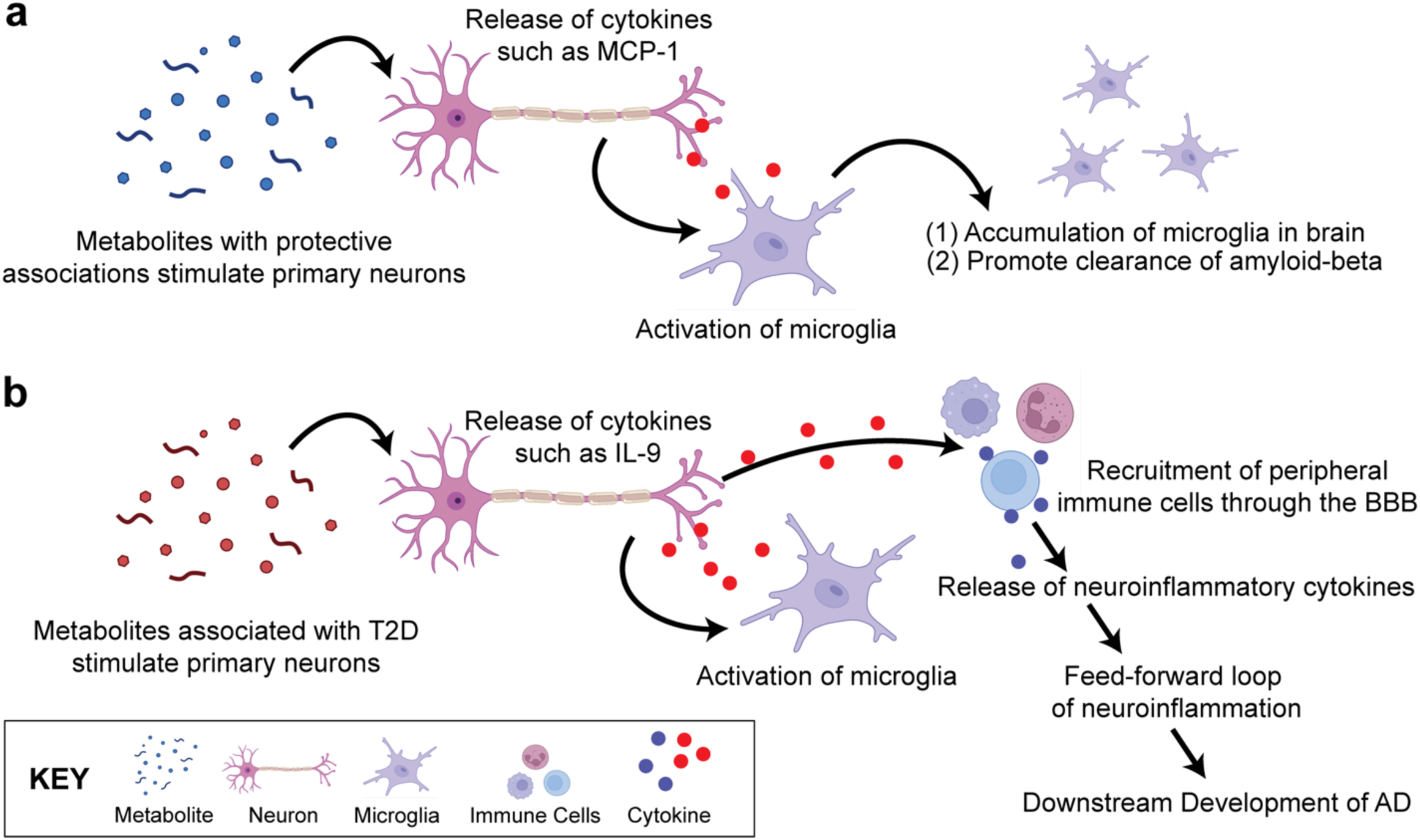
Hypothesized Pathway of the Shared Neuroinflammatory Response. **(a)** In a healthy state, few metabolites may cross the BBB through specialized transport. The stimulated neurons produce cytokines that may activate glia such as microglia, which will accumulate in the brain and promote clearance of amyloid-beta and other debris in the central nervous system. **(b)** In a diseased state, metabolites may cross the BBB in higher concentrations, and stimulate the neurons and glia cells. This chronic, low-grade stimulation of neuronal cells may result in the eventual breakdown of the BBB, leading to downstream migration of immune cells and more metabolites to enter the brain. The release of neuroinflammatory cytokines may generate a feed-forward loop of neuroinflammation, potentially leading to the development of eventual AD. (Created with BioRender.com).

A limitation of our work is that the neuron culture only represents one of the many cells located in the central nervous system in a single-direction pathway where neurons respond to different disease-associated metabolites. Cytokines are complex signaling molecules that themselves cannot be simply categorically into simple associations. Likewise, it is difficult to fully classify a metabolite to be associated with a specific disease, as human biology is multi-dimensional. In the future, preparing an experiment with different co-culture models may pose alternative methods to study signaling networks related to T2D as a risk factor for AD development.

Our work examined the patterns of neuroinflammatory cytokines released by primary mouse neurons when stimulated by metabolites differentially produced in AD and T2D. Our findings show that metabolites with similar disease associations result in similar profiles of differentiating cytokines such as MCP-1. Understanding the patterns of cytokines released by neurons will allow us to infer the potential neighboring cells that may be activated. Our results suggest a need for further studies to investigate T2D-driven neuroinflammation as a contributor to AD.

## MATERIALS AND METHODS

### Candidate Metabolite Selection

The selection criteria for inclusion of metabolites in our study were (1) that the metabolite be differentially abundant in AD, T2D, or both, (2) that the metabolite be BBB-permeable, and (3) that two or more studies supported these associations. We used the search terms “metabolites present in Alzheimer’s disease” and “metabolites present in type 2 diabetes” in Google Scholar and PubMed. The identified metabolites were then given an association with detrimental or protective characteristics based on findings from at least two independently published literature. The associations were established by using search terms “[identified metabolite name] and Alzheimer’s disease” and “[identified metabolite name] and type 2 diabetes.” In our search criteria, while there is an increased BBB permeability shared in both T2D and AD, all metabolites were verified to have the ability to cross the blood-brain barrier based on specialized transport mechanisms or favorable chemical properties. We confirmed this through literature findings with the keyword search “[metabolite name] and BBB permeability”^62–70^. With at least two studies supporting the associations, we identified nine candidate metabolites for follow-up studies, which include lauric acid, asparagine, fructose, arachidonic acid, aminoadipic acid, D-sorbitol, retinol, L-tryptophan, and niacinamide for the stimulation of primary culture of neurons (**Table 1**).

**Table 1.** Metabolites selected from literature for neuron stimulation.

| Metabolite | Associations | References |
| --- | --- | --- |
| Lauric Acid | T2D<sup>↑</sup> | 55,83 |
| Asparagine | T2D<sup>↑</sup> | 56,84 |
| Fructose-6-Phosphate | AD<sup>↑</sup>, T2D<sup>↑</sup> | 85,86 |
| Arachidonic Acid | AD<sup>↑</sup>, T2D<sup>↑</sup> | 87,88 |
| Aminoadipic Acid | AD<sup>↑</sup>, T2D<sup>↑</sup> | 84,89 |
| D-Sorbitol | AD<sup>↑</sup>, T2D<sup>↑</sup> | 84,90 |
| Retinol (Vitamin A) | AD<sup>↓</sup>, T2D<sup>↓</sup> | 83,91,92 |
| L-Tryptophan | AD<sup>↓</sup> | 93,94 |
| Niacinamide | AD<sup>↓</sup> | 84,95 |

### Primary Neuron Culture

All animal procedures were performed in strict accordance with the guidelines approved by the Penn State College of Medicine Institutional Animal Care and Use Committee (PROTO201800449). The primary neurons used in this investigation are derived from embryonic litters from pregnant CD-1 mice (Charles River Laboratories, strain 022). Two pregnant mice (gestational day 17) were sacrificed by decapitation using a guillotine. Between the two pregnant mice, a total of 25 embryonic pups were sacrificed by decapitation using surgical scissors. The isolated brains were immediately placed in cold HEPES-buffered Hank’s Balanced Salt solution for dissection. After the meninges were removed, the cortical cap of the brain was isolated.

Primary neuron culture was performed according to validated methods^71^. The isolated cortices were transferred to a conical tube containing warm embryonic plating medium: Neurobasal Plus (Gibco), 10% fetal bovine serum (Gibco), 1x GlutaMAX (Gibco), 1x Penicillin-Streptomycin (10,000 U/mL, Gibco). Cortices were manually triturated using a pipette in the embryonic medium, and the overall cell concentration was determined by the Countess II automated cell counter (Invitrogen). The 6-well plates coated with 0.1 mg/mL Poly-D-Lysine were plated with the cell suspension at a density of 5,196,500 cells/well. After 24 hours, the embryonic media was replaced with neuronal media: Neurobasal Plus (Gibco), 1x B27 Plus supplement (Gibco), 1x GlutaMAX (Gibco), 1x Penicillin-Streptomycin (10,000 U/mL, Gibco). Half of the neuronal medium was replaced after four days. After inspecting the plates for confluency, the neuronal medium was aspirated, and neurons were stimulated with metabolites dissolved in media on day 8 with a final concentration of 300 uM. On day 11, cell media was collected and snap-frozen in liquid nitrogen for future cytokine quantification with the Luminex system.

### Established Metabolite Stimulation Concentration

Metabolite treatment concentration was determined by dose-response neuron viability (Invitrogen, catalog no. L3224) in the nano and micromolar range. We established the range we tested for our live/dead assay based on the ranges tested in these metabolites with different cell lines in prior studies^72–80^. Like the primary neuron culturing methods, one pregnant CD1 mouse at gestational day 15 (n=12 embryos) was sacrificed and plated, with the adjustment of 175,000 cells/well in a 96-well plate. On day 8, we stimulated neurons with twelve logarithmically increasing concentrations (1 nM, 3 nM, 10 nM, 10 nM, 30 nM, 100 nM, 300 nM, 1 uM, 3 uM, 10 uM, 30 uM, 100 uM, and 300 uM) (**Supplementary Fig. S2**). Each metabolite was first dissolved in dimethyl sulfoxide, water, or phosphate-buffered saline before being mixed with neuronal media. The vehicle controls (no metabolite added) matched the solvent composition respective to each metabolite group. The dimethyl sulfoxide concentration did not exceed 0.3% to ensure cell viability. On day 11, we assayed neuron viability using live/dead staining (calcein acetoxymethyl and ethidium homodimer, catalog no. L3224, Invitrogen). Fluorescence was measured using the Spectramax i3x microplate reader (Molecular Devices). We found no significant decline in cell viability as the concentration increased, thus, the 300 uM concentration was selected as the established concentration for our assays, with the rationale that high acute exposure will exhibit a similar response to low chronic exposure.

### Cytokine Quantification

Samples, blanks, and quality controls were loaded into a 384-well plate in technical triplicate, and cytokine concentrations were quantified using the Luminex Bio-Plex 3D platform. The assay was carried out with the Bio-Rad BioPlex 23-Plex Pro Mouse Cytokine Kit, which contains a panel to quantify pro- and anti-inflammatory cytokines (Eotaxin, G-CSF, IFN-γ, IL-17A, IL-1β, IL-1α, IL-10, IL-12p40, IL-12p70, IL-13, IL-2, IL-3, IL-4, IL-5, IL-6, IL-9, KC, MCP-1, MIP-1β, MIP-1α, RANTES, and TNF-α) from Bio-Rad (Catalog no. M60009RDPD). The manufacturer’s protocol was modified to accommodate a 384-well format by adding magnetic beads and antibodies at a reduced volume^81,82^. On the day of the assay, the collected samples were thawed on ice. All steps that involved plate washing were performed with the Hydrospeed plate washer with magnets (Tecan). The Luminex instrument calibration and verification were conducted on the day of the assay.

### Data Pre-Processing and Normalization

Cytokines with 25% or more of the samples reading below the lower limit of quantification were excluded from the analysis. The remaining 15 quantified cytokines were retained for downstream analysis. To prepare the data for analysis, any individual values below the lower limit of quantification were replaced with the lowest respective standard value (2.1% of data). Cytokine concentrations were then normalized by a log_2_ ratio of the individual measurement divided by the average of the triplicate vehicle concentrations. All data analysis was conducted in RStudio (version 1.4.1717 Juliet Rose).

### Unsupervised Hierarchical Clustering and Principal Component Analysis

The log_2_ normalized data was used for unsupervised hierarchical clustering and principal component analysis (PCA). For a global comparison across different metabolite groups and cytokines, a heatmap was generated using the *pheatmap* package in RStudio (package version 1.0.12). The *factoextra* package was used for PCA, with the input data scaled and normalized to the vehicle groups (package version 1.0.7).

### Statistical Analysis

To determine the statistical significance of differences in cytokine levels across metabolite treatment groups, we performed a non-parametric Kruskal-Wallis test on the data. A Benjamini-Hochberg was performed on the Kruskal-Wallis test to correct the false discovery rate (FDR) of multiple comparisons. Significance was determined if an FDR *q* value was less than 0.10. We performed post-hoc testing with a non-parametric Mann-Whitney test on cytokines identified to be significantly differing across the four groups of disease-associated metabolites (AD-protective, AD/T2D-protective, T2D, and AD/T2D), with a *p* value less than 0.05 considered significant. Significantly differentially abundant cytokines were visualized with box and whisker plots.

### Partial Least Squares Discriminant Analysis

We performed partial least squares discriminant analysis (PLS-DA), a multivariate dimensionality-reduction technique in RStudio using *mixOmics* (package version 6.16.3). For PLS-DA, the normalized log_2_ fold change of cytokine concentrations secreted by primary neurons were independent predictor variables and our dependent variable was the disease association of the metabolites. We constructed three PLS-DA models using cytokine profiles to predict (1) 4-way discrimination between all metabolite groups (AD-protective, AD/T2D-protective, T2D, and AD/T2D), (2) AD-protective vs. AD-associated, and (3) T2D-protective vs. T2D-associated metabolite stimulation conditions. The number of latent variables (LV) selected for each model was determined by a three-fold cross-validation repeated randomly one hundred times based on the model with the lowest cross-validation error rate.

In our PLS-DA model, we identified the most important cytokines contributing to the model’s overall predictive accuracy by the variable importance in projection (VIP) score. The VIP scores were calculated by using the *mixOmics* package, which represents the strength of contribution from each cytokine to the results of the PLS-DA model:

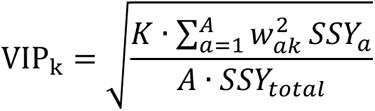

Where *K* is the total number of cytokine predictors, *A* is the number of PLS-DA components, *w_ak_* is the weight of predictor *j* in the *a*^th^ LV component, and *SSY_a_* is the sum of squares of the explained variance for the *a*^th^ LV component. The *SSY_total_* represents the total sum of squares explained in all of the LV components. Since the VIP accounts for normalization, a score greater than 1 indicates an important variable within the model.

## ABBREVIATIONS

AD: Alzheimer’s disease
BBB: Blood-brain barrier
FDR: False discovery rate
G-CSF: Granulocyte colony-stimulating factor
IFN-γ: Interferon-γ
IL: Interleukin
KC: Keratinocyte chemoattractant
LV: Latent variable
MCP-1: Monocyte chemoattractant protein-1
MIP-1α: Macrophage inflammatory protein-1α
MIP-1β: Macrophage inflammatory protein-1β
PCA: Principal component analysis
PLS-DA: Partial least squares discriminant analysis
RANTES: Regulated on activation, normal T cell expressed and secretion
T2D: Type 2 diabetes
TNF-α: Tumor necrosis factor-α
VIP: Variable importance in projection

## ACKNOWLEDGEMENTS

This work is supported by an award from the Good Ventures Foundation and Open Philanthropy, as well as start-up funds from Purdue University Weldon School of Biomedical Engineering (DKB and BKB). This work is also supported by R21AG068532 from the National Institute on Aging (EAP). BKB is supported by the NIH T32 predoctoral fellowship T32DK101001 from the National Institute of Diabetes and Digestive and Kidney Diseases. BKB acknowledges the National Science Foundation for support under the Graduate Research Fellowship program (GRFP) under grant number DGE-1842166. MKK is supported by training fellowship T32NS115667 from the National Institute of Neurological Disorders and Stroke. RMF is supported by NIH NRSA predoctoral fellowship F31AG071131 from the National Institute on Aging. The authors thank Javier Muñoz Briones (Purdue University) for support in the Luminex assay. The authors also thank Raymond Krajci (Case Western Reserve University) for verifying reproducibility of the code used for analysis.

## AUTHOR CONTRIBUTIONS

BKB: Conceptualization, data curation, formal analysis, funding acquisition, investigation, methodology, visualization, writing-original draft, writing-review & editing. MKK: Formal analysis, investigation, methodology, writing-review & editing. RMF: Formal analysis, investigation, methodology, writing-review & editing. EAP: Conceptualization, funding acquisition, methodology, project administration, resources, writing-review & editing. DKB: Conceptualization, funding acquisition, methodology, project administration, resources, writing-review & editing.

## DATA AVAILABILITY

Generated data for analysis is included and available in the Supplementary Files of this article.

## CODE AVAILABILITY

All code is publicly available at https://github.com/Brubaker-Lab/AD-T2D-Cytokine-Manuscript.

## SUPPLEMENTARY FIGURES

**Supplementary Figure S1.**
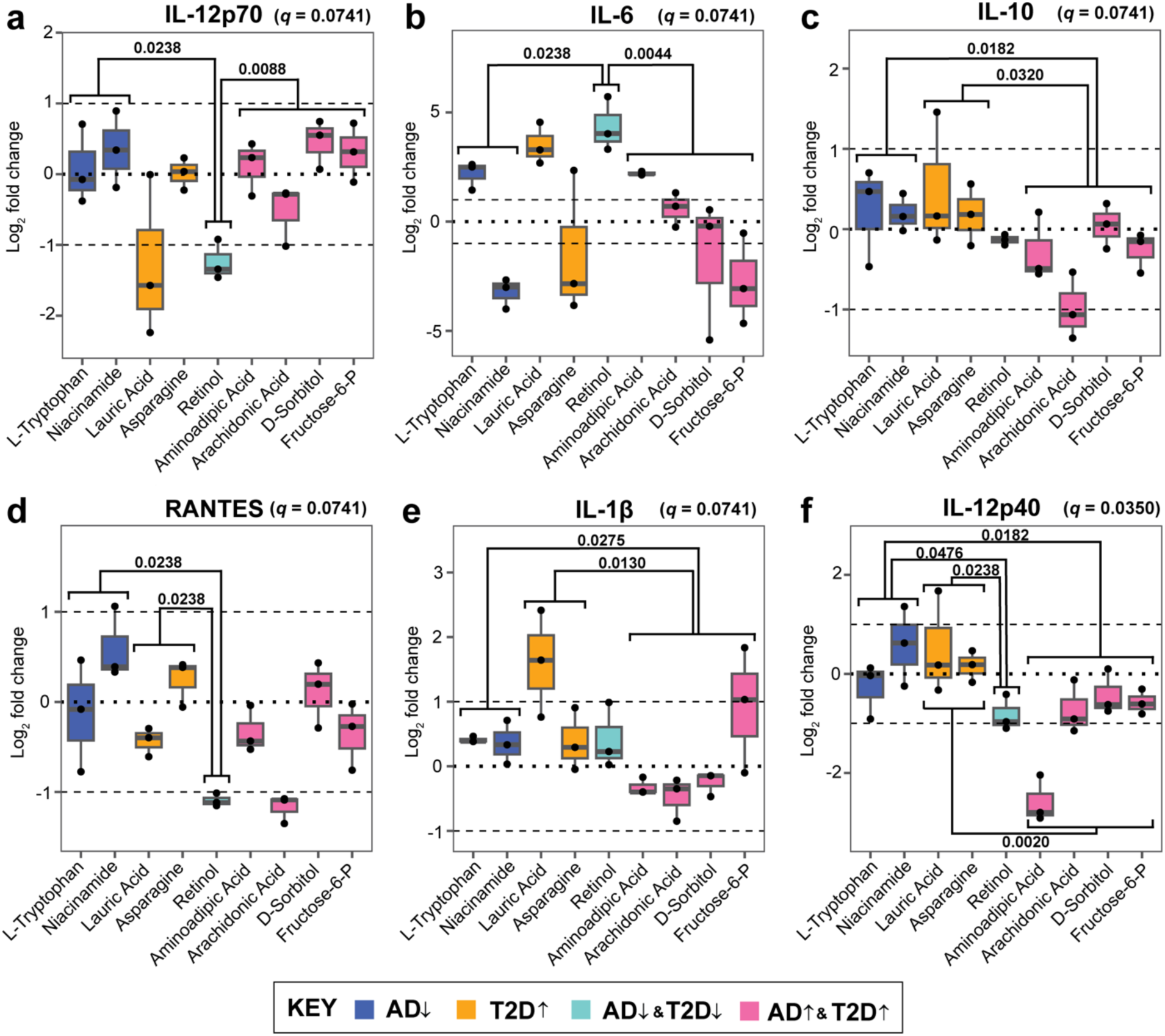
Additional Significantly Reported Results on Neurons treated with different disease associations. The log_2_ ratio of cytokine concentration to vehicles that were determined significant from a Kruskal-Wallis test (FDR *q* value next to the cytokine). Mann-Whitney pair-wise testing was applied to each metabolite group based on disease association (significance denoted within the plot). The cytokines include **(a)** IL-12p70, **(b)** IL-6, **(c)** IL-10, **(d)** RANTES, **(e)** IL-1β, and **(f)** IL-12p40.

**Supplementary Figure S2.**
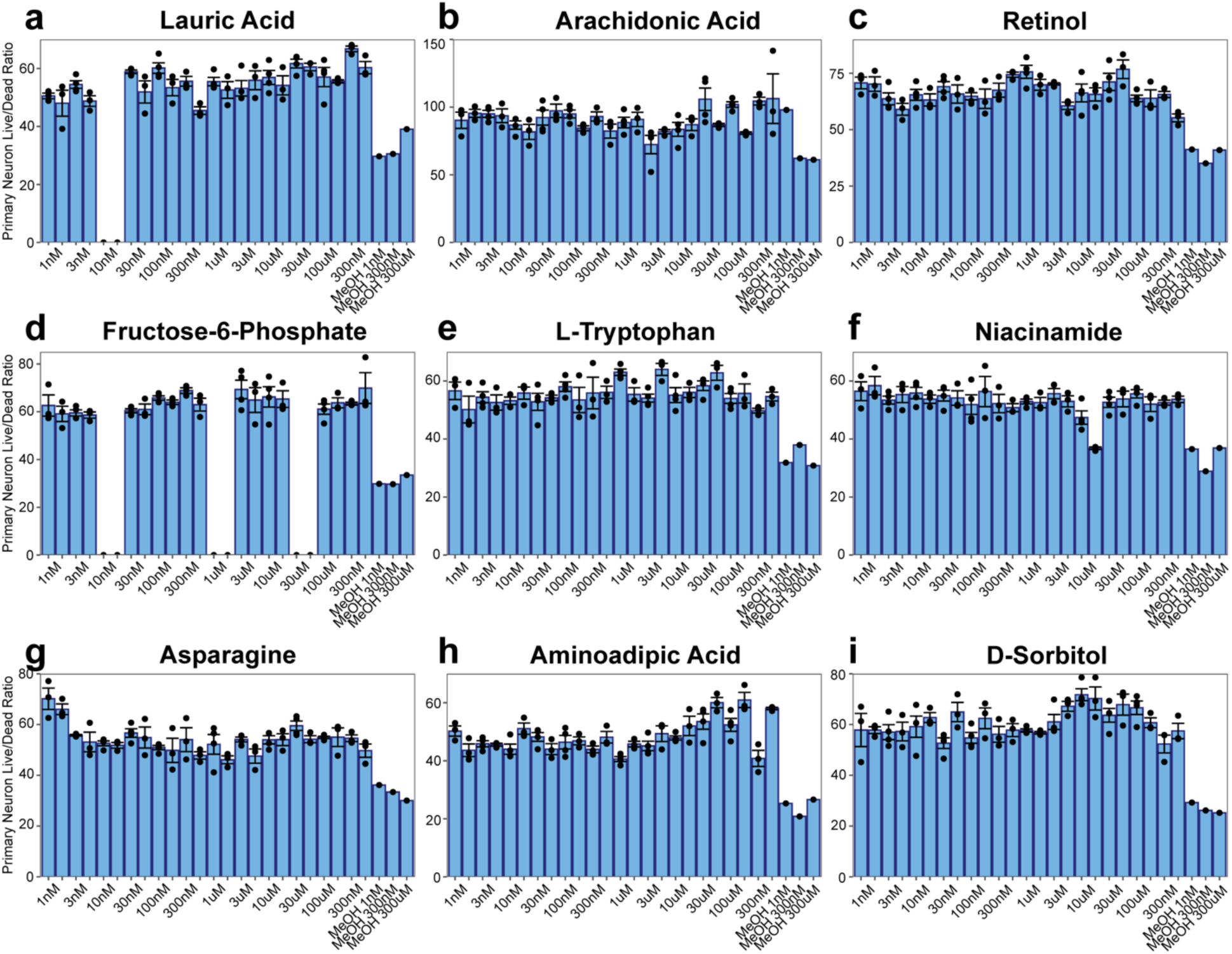
Live-dead ratio of each metabolite stimulation on primary neuron culture. Live-dead assay was performed on **(a)** lauric acid, **(b)** arachidonic acid, **(c)** retinol, **(d)** fructose-6-phosphate, **(e)** L-tryptophan, **(f)** niacinamide, **(g)** asparagine, **(h)** aminoadipic acid, and **(i)** D-sorbitol. Per each concentration, the metabolite (left-side bar) and vehicle (right-side bar) is reported. Missing data is due to extreme outlier (lauric acid) and limitation of primary neurons (fructose-6-phosphate). Live-dead ratio is displayed as the mean ± standard error of the mean.

